# Optimal division asymmetry in *E. coli* bacteria

**DOI:** 10.1101/2025.04.10.648155

**Authors:** Ulrich K. Steiner, Audrey Proenca, Murat Tugrul

## Abstract

Cell division asymmetry underlies many fundamental biological processes, including the evolution of aging. Despite this importance, understanding whether the observed division asymmetry has been optimized by evolution remains little explored. Here we first show that size and transcription signal asymmetries have evolved to an optimum, as increased or decreased asymmetries lowers population growth. We then highlight how variance and covariance in growth and transcription signal asymmetry influence population growth. This latter exploration reveals how sensitive deviations to the optimum are, and that selective forces act primarily on variance in transcription signal asymmetry and less so on other cell intrinsic noise or growth asymmetries. Our findings are based on advancing integral projection models to two traits and we parameterize these models with single-cell data of mother and daughter *E. coli* bacterial cells collected using microfluidic devices. We discuss how trade-offs between growth and the transcription signal we explore has shaped this optimum. Our findings provide insights on how division asymmetries and resulting cellular heterogeneity can have profound biological consequences—contributing to processes such as cancer development, antibiotic treatment failure, and the rejuvenation of cell lineages, including in mammalian stem cells.

## Introduction

Most bacterial cells reproduce by binary fission. Contrasting classical assumptions of symmetric fission (Williams 1957; Hamilton 1966; Partridge & Barton 1993), the two emerging offspring cells are not identical (Stewart *et al*. 2005; Steiner 2021). Such asymmetry is crucial to understand the evolution of aging, the division of labour, and the origin of multicellularity. If one assumes that perfect symmetric division could be achieved, cells would either be non-aging and potentially immortal, or cell populations would constantly age by accumulating some aging factor, and this accumulation would overwhelm the population leading to its final crash (Ackermann *et al*. 2007; Pen & Flatt 2021; De Vries *et al*. 2023). In multicellular organisms, asymmetric cell division facilitates the generation of diverse cell types, and fields such as developmental biology focus on differentiation and asymmetry originating from cell division, including mammalian stem cells (Neumüller & Knoblich 2009). In unicellular, basal prokaryotic cells, such as bacteria or archaea, cell type differentiation is often lacking or hard to evaluate. Discussions linger whether asymmetry is needed for any cell division processes to occur and if so, whether the observed asymmetry is optimal.

Empirical evidence accumulates that also in procaryotic cells, perfect symmetric fission is challenging to achieve. The asymmetry between the two emerging offspring cells manifests in differences in the membrane age, as most bacteria synthesize a new cell membrane at the central plane when dividing. This synthesis at the cell’s central plane leads to an old pole side and a new pole side of a cell. The age of the pole allows to distinguish, over multiple divisions, a mother (old pole) cell and a daughter (new pole) cell (Stewart *et al*. 2005). Mother and daughter cells differ in growth rates and division rates, with mothers showing reduced growth and division (1–6). Physiological asymmetries are not only revealed by growth differences and by aggregates of (misfolded-)proteins that are located at the old pole of the mother cell (1, 6), but also when dispersed transcriptional signal differs in their intensities among mother and daughter cells (5, 7). Asymmetries not only arise through the partition of cytoplasm content but also through unequal physiological activity among mother and daughter cells (Proenca *et al*. 2024), which become progressively apparent with increasing (mother-pole) age (1, 5–7). Recently, even the long-assumed morphologically symmetric division of bacteria such as *E. coli* has been challenged, by showing that mothers increase in size with age, while daughters conserve their size at divisions (6, 8). The importance of such asymmetry is also revealed in survival with mother cells exhibiting higher mortality compared to daughter cells (Steiner *et al*. 2019).

Theoretical explorations illustrate that division asymmetry should be optimized to maximize population growth, i.e., i.e. fitness (9, 10). Predicting the optimum of asymmetry has been challenging, as empirical quantification of asymmetries is rare and unconnected to fitness. Many models indicate intermediate optimal asymmetries, though perfect symmetry could also arise when the accumulation of damage can be prevented by growth, i.e. biomass production, that dilutes damage production at a similar rate (Clegg *et al*. 2014). Some models that explain how aging can evolve under the assumption of perfect division symmetries (Pen & Flatt 2021) were questioned as they included hidden among cell asymmetries (De Vries *et al*. 2023). Combined, the theoretical predictions on the outcome of asymmetry depends on underlying model assumptions, here we do not evaluate these assumptions but explore whether observed division asymmetries are at an optimum.

We ask this question of an evolved optimum by quantifying mother-daughter pair division asymmetries of single-cell *E. coli* bacteria and link these single-cell asymmetries to population growth. We use size and transcriptional signal of a general stress response regulator (*rpoS*) that regulates about 10% of all *E. coli* genes. This rpoS transcription factor is negatively correlated with growth (5, 11). Structured population models (Integral projection models, IPM) that use continuous traits associated with fitness, such as size, have been frequently applied in demography and population biology to link individual to population dynamics(12, 13), but such models have been less applied for single-cell bacteria studies, where size is closely linked to reproduction and the major gene expression regulator rpoS forming proxies for individual fitness. Here, we expand previous IPM’s to allow for full interaction and projection of two continuous traits: in our study, size and transcription signal. Using these expanded models, we evaluate whether altered asymmetries at cell division would lead to increased population growth. In addition to exploring mean changes in asymmetry, we also evaluate how variance and covariance around division asymmetries, and in cell intrinsic stochastic processes, affect population growth. Such exploration of altered variance is easily achieved by these structured population models as they are inherently stochastic, being rooted within Markov chain theories. Exploring the sensitivity to perturbations by evaluating variance in asymmetry, growth, and gene regulation, provides an understanding of whether these factors are under selection, and of the selective strength acting on them. Increased variance in vital rates should reduce population growth compared to mean constant vital rates and hence selection should act against variability (Tuljapurkar 1982), though empirical evidence suggest only weak selection on stochastic variability in life courses (Steiner & Tuljapurkar 2023). Other theoretical models, focusing on variability in generation time, suggest positive effects of variability on population growth, indicating selection favouring variability (Hashimoto *et al*. 2016; Genthon 2024). Our exploration on variance in growth, transcription signal, and asymmetry will help to evaluate the effect and strength of selective forces on such stochastic processes and cell-intrinsic noise that generate cell-to-cell heterogeneity.

In the following we first expand integral projection matrix models to two traits. We then report on our findings parameterizing these models with data from single-cell growth and transcription signal dynamics. Finally, we discuss how quantifying asymmetries in bacteria helps to understand aging, the generating processes of cell-to-cell heterogeneity, and the consequences of asymmetries and individual heterogeneity that go beyond aging in bacteria.

### Model

We advance integral projection models (IPM) that classically integrate one continuous variable (Ellner & Rees 2006), to a model that integrates two continuous variables, here size and transcription signal (*rpoS*). Previous versions of IPM’s described how to integrate across two continuous variables, but in these models, only one of the two variables is directly projected, and the other one could be seen as a covariate (Ellner *et al*. 2016). We categorize projection kernels and describe our approach below.

### Classical IPM

We first consider a classical IPM with one variable as

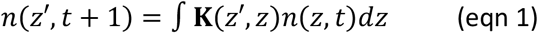

where *n*(*z*′, *t* + 1) describes the population vector of individuals with trait characteristic *z*′at time *t* + 1. This population vector is computed by the integral of the kernel, with the integral spanning all possible range of trait z (*dz*). The core part of the kernel, **K**(*z*′, *z*), describes the relationship between trait *z* individuals at time *t* to the trait *z’* individuals at time *t+1*. In most applications of IPMs in ecology and demography, the trait that is tracked is size, as it is conveniently measured and related to fitness. The kernel **K**(*z*′, *z*) is often split into a survival and growth kernel **P**(*z*′, *z*) and a fecundity kernelF(*z*′, *z*), i.e.

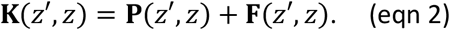

The survival and growth kernels express the probability that an individual survives from time *t* to time *t+1*, and if the individual survives, the probability of a trait *z* individual at time *t* to become a trait *z’* individual at *t+1*. This **P**(*z*′, *z*) part of the kernel can hence be decomposed into the survival probability *s*(*z*) and the trait transition probability, **G**(*z*′, *z*); for sizes this would be growth of size *z* individuals at time *t* to size *z’* individuals at time *t+1*, so

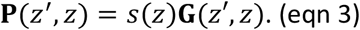

The fecundity kernel expresses how many type *z’* offspring being produced at time *t+1*, by individuals of trait *z* at time *t*. This part includes the binary probability, *p*_*b*_(*z*), the yes/no probability of a size *z* individual at time *t* to reproduce within the time interval from *t* to *t+1*. The number of offspring that are produced in the time interval *t* to *t+1, p*_*r*_*b*(*z*). Note, for binary fissioning bacteria only one offspring is produced, thus, *p*_*r*_*b*(*z*) = 1. We therefore omit *p*_*r*_*b*(*z*) when expanding below to two continuous traits, but it could easily be included for any more generalized case. Finally, the **C**_1_(*z*^′^, *z*), describes the trait distribution of offspring produced by a size *z* individual. Combined, the fecundity kernel can hence be written as

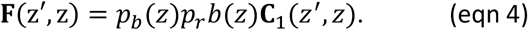

### Advancing IPM’s to two continuous variables

To advance a one variable IPM (eqn 1) to a two variable IPM we simply need to expand the population vector and kernel to incorporate a second variable,

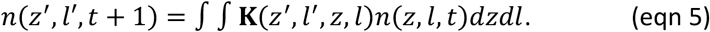

In the following we consider the two variables as the size at time t (z) and size at time t+1 (z’) and the transcription factor (TF) signal at time t (l) and TF at time t+1 (l’) respectively. The decomposition in survival-growth-TF and fertility parts

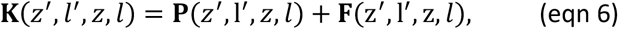

follow accordingly, so can these be decomposed into

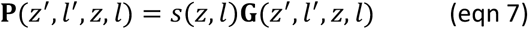

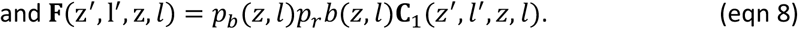

All probabilities, transitions, and trait distributions of the offspring, now just depend on size (z) and TF level (l) at time *t* of mother cells.

For estimating Kernel parameters, we refer here to the method description (see also Fig. 2, and Supplemental Figures S2 & S3). We used the estimated **K**(*z*^′^, *l*^′^, *z, l*) = **P**(*z*^′^, l^′^, *z, l*) + **F**(*z*^′^, l^′^, z, *l*) to project the population vector of *n*(*z, l, t*) to *n*(*z*^′^, *l*′, *t* + 1) starting with a uniformly distributed *n*(*z, l*) vector at time *t* = 1 (Fig. S3). This projection illustrates the convergence time to reach stable stage distribution and reveals the population growth, i.e. fitness (Fig. 3a&b).

**Fig. 1:**
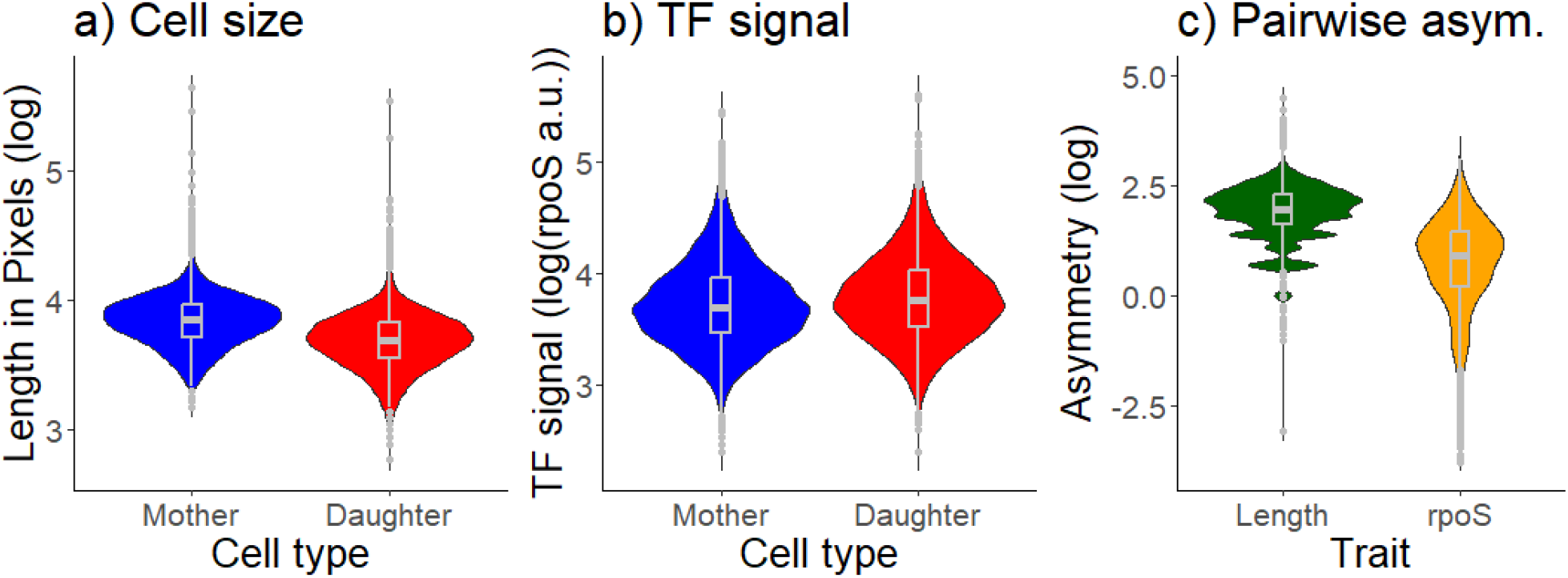
Size (a) and TF, transcription signal (b) between mother and daughter cells right after cell division. Cell types differed in length and in TF signal. Statistical testing was done using model comparisons between an intercept only model and a model that accounts for cell type difference (Cell size: ΔAIC2141; TF signal: ΔAIC 83). c) The pairwise difference between mother-daughters highlights the size and transcription asymmetry right after fissioning.

**Fig. 2:**
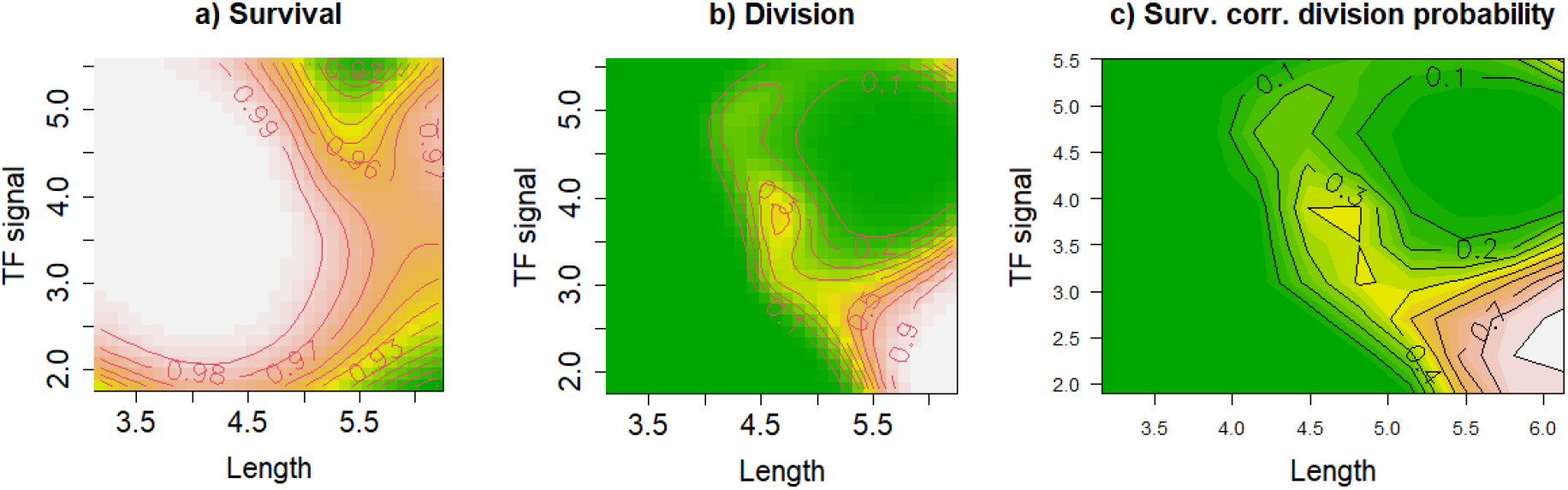
a) Survival probabilities (per 4 Min), *s*(*z, l*), b) division probabilities (per 4 Min), *p*_*b*_(*z, l*), for mother cell length (log transformed pixels) and TF signal (log transformed a.u.). Higher survival and division probabilities are lighter colours, lower survival and division probabilities are darker green coloured (see numbers on isolines). c) Survival corrected reproduction probabilities, *s*(*z, l*)*p*_*b*_(*z, l*) for mother cell length (log transformed pixels) and TF (log transformed a.u.) signal. Details of parametrization and estimation using GAMs are given in the methods.

**Fig. 3:**
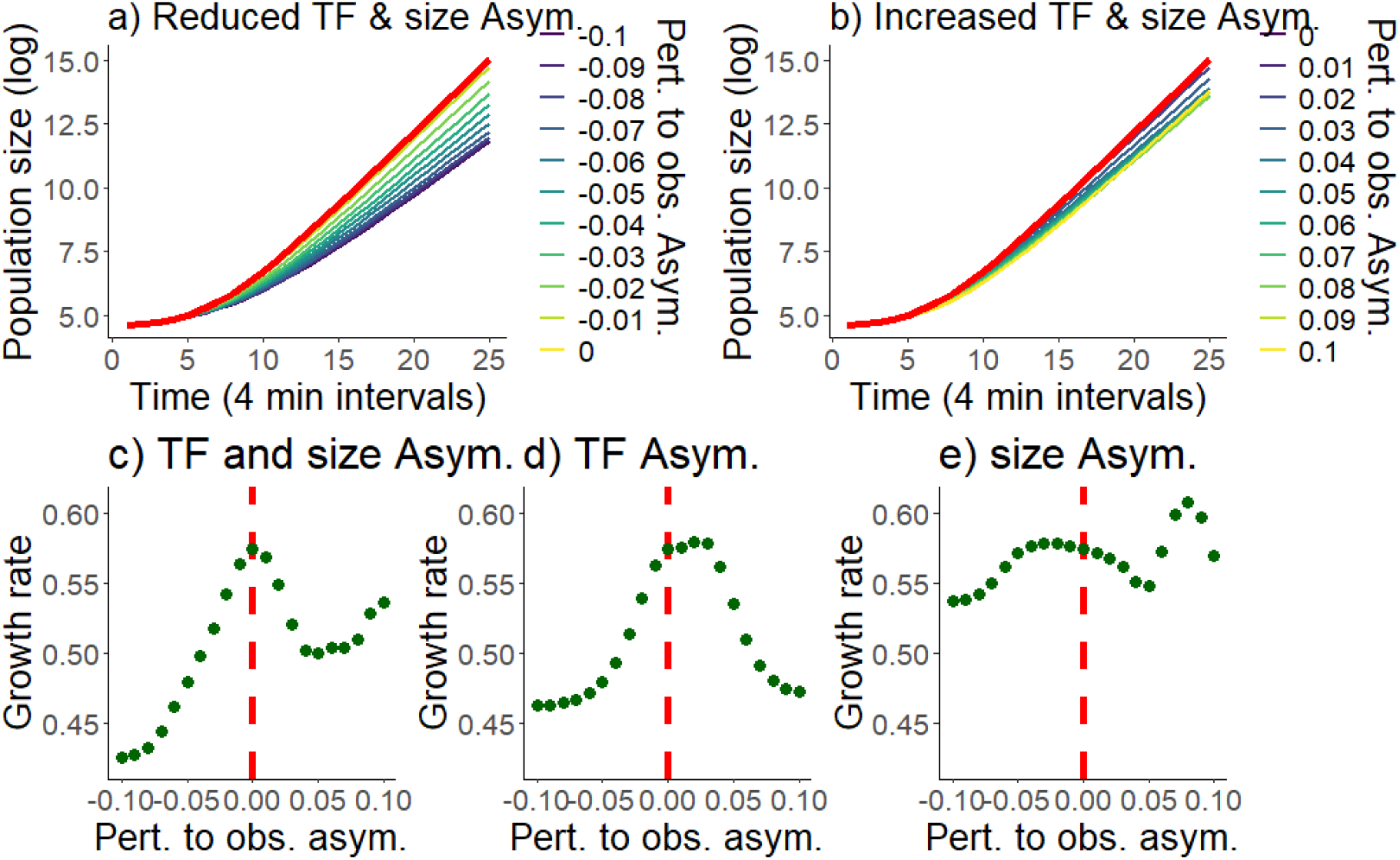
Simulations of a) Population growth (log Population size) over time (4 min intervals) for (a) reduced and (b) increased asymmetries, compared to observed asymmetries, in TF and cell size after division. The red line marks the observed, i.e. unperturbed, asymmetry. c-e) Exponential growth rates for different perturbation to the observed asymmetries when (c) TF and size asymmetries are perturbed, when (d) only TF asymmetries are perturbed and size is fixed at the observed asymmetry, or when (d) only size asymmetries are perturbed and TF is fixed at the observed asymmetry. c) summarizes the data from (a & b) with the data points left of the hatched red line illustrating growth rates shown in (a), and data points to the right of the hatched red line illustrating growth rates shown in (b). The hatched line in c-e) marks the observed asymmetry.

### Exploring asymmetry and variance effects on fitness

We start by parameterizing the model with the observed asymmetry and variance (see Method), this computes the population growth and thereby fitness. We then alter this initial model by numerically increasing or reducing asymmetry, variance, and covariance for size and TF at cell division, and estimate the population growth for these perturbed models. The optimal asymmetry is then simply the asymmetry that results in the largest population growth. Perturbation or sensitivity analyses are frequently used to evaluate optima and to quantify selective forces acting against perturbations, as illustrated by demographic buffering or sensitivity arguments (14, 15). The reasoning is, that a small deviation from the optimum, that causes a large fitness reduction, should be strongly selected against, while a larger deviation from the optimum, that has only small fitness effects, should not be strongly selected against (14, 15). We apply this reasoning to assess the selective forces acting on variance in asymmetry. Similar as for mean division asymmetry in growth and rpoS between mother and daughter cells, and the variance in division asymmetry, we can also quantify the effect of variance and covariance on fitness within cells. That is, we increase or decrease rpoS or growth noise, stochasticity, or uncertainty of cells that are not dividing, and estimate the effect of such increased noise for fitness. Larger sensitivity to increased noise should again indicate stronger selective forces acting against such noise.

## Results

Before we investigate whether there is an optimal asymmetry by modelling perturbed asymmetries, we report on the observed asymmetries that arise directly from the empirical data and illustrate some of the underlying patterns of the model parameterized with these observed data (5). Size asymmetry between mother and daughter cells reveals that mothers being about 17% larger than daughters right after division, and the fluorescence asymmetry is shown by mothers having about 5% lower transcription signal compared to daughter cells (Fig. 1). Just before division, mothers mean size was 4.33 μm (86.9 pixels; log = 4.46; 100 pixel = 4.89 μm) and TF signal strength was 45.3 a.u. (log 3.81); right after division, mothers were 2.32 μm (47.4 pixels; log = 3.86) long and TF signal strength was 44.7 a.u. (log = 3.80). Their daughter cells showed mean sizes of 1.98 μm (40.6 pixels; log = 3.7) and TF signal strength of 47.2 (log = 3.85). Hence, mean size asymmetry after division is ∼1.17 (2.32/1.98), with mother cells being larger than daughter cells, and mean fluorescence asymmetry is ∼0.95 (44.7/47.2), with daughter cells showing higher TF signal compared to mother cells. For these and following analyses we used 95272 size and TF transition events over the 36 hour-lasting experiment. In addition we used 7678 division events for which the TF and length of the mother at time *t* and the TF and length of the daughter at time *t+1* was known. In most cases, TF and length measures have been log-transformed for better visualization and to better fit the expectation of normally distributed errors.

Survival probabilities are lower at large cell sizes, in addition, TF levels mainly influence survival of larger cells with high or very low TF signal (Fig. 2a). Division probabilities are highest for large cells with low TF signal. Small cells do not divide, and high TF signal is known to correlate with low cell elongation rates (16), which can explain why large cells with high TF (low growth) do not divide frequently (Fig. 2b). Combined, overall survival has been high, particularly for small cells, showing that mainly larger cells risk death, but these are also the cells that divide and thereby reproduce (Fig. 2c).

### Optimal asymmetry

Our perturbation analysis highlights, that the observed mother daughter asymmetry at division in length and TF signal results in the fastest population growth and reveals the highest fitness (Fig. 3a & 3b red line). Reducing or increasing the asymmetry always leads to lower fitness (Fig. 3a & b, see optimum in Fig. 3c). Interestingly, investigating only perturbed TF asymmetry (Fig. 3d) or only size asymmetry (Fig. 3e) suggests that a sightly higher TF asymmetry and size asymmetry should result in an increased growth rate. For size, the highest fitness (Fig. 3e) would be achieved by perturbation of ∼3%, and for the TF asymmetry an perturbation of ∼2% would lead to the highest expected fitness. Note, as mother cells are larger than daughter cells, but mother cells have lower TF signal than daughter cells (Fig. 1) increased asymmetry is achieved by a perturbation of negative value for size (Fig. 3e), and perturbation of positive value for TF (Fig. 3d). However, the combination of perturbation as explored in Fig. 3c that increases or decreases asymmetry in both variables, does not lead to increased fitness (Fig. 3c). Likely trade-offs between growth and TF, that are known to negatively correlate the two variables, balance a higher or lower asymmetry(16). The observed asymmetry (red lines) seems to be the optimal solution selection has acted on. Further, notice that TF asymmetries seem to influence population growth rates to a larger degree than size asymmetries as the population growth is more strongly affected by TF asymmetry manipulations than length asymmetry manipulations (Fig. 3d&e). This latter interpretation needs to be taken with caution, as we are altering asymmetries by multiplying in a range of values (-0.1 to 0.1) partly on measures with arbitrary units (TF). Therefore, the strength of altered asymmetry might not be directly compared between the two variables (length and TF).

### Sensitivity of the optimal asymmetry and intracellular noise

Aside of the fitness effects of mean asymmetries at division we explore in Fig. 3, we also explore increased and decreased variance or noise in growth, TF, and division asymmetry and how these altered variances influence population growth, i.e. fitness (Fig. 4). In doing so, we explore how sensitive the detected optimum in asymmetry is (Fig. 3), or in different words, how strong deviations, of for instance among individual cell variation, around that optimum influence fitness. The higher the sensitivity, the stronger selection should act against stochasticity—or noise—in size or transcription signal asymmetry, intracellular growth noise, and transcription factor noise. Stochastic population theory predicts that reduced temporal variability in vital rates increases population growth (17, 18), theories on demographic buffering suggest that selection should act the strongest on matrix parameters that increase variability, that is the most sensitive matrix parameters should be selected more strongly against variance than less sensitive parameters (14), and fundamental evolutionary theories suggest that variability among genotypes should be selected against in a given environment (19). Most previous studies explored variance (stochastic) effects of one variable, but here we explore two interacting variables. We therefore evaluate both the effect of variance and covariance in size and transcription factor signal on population growth. Aligned with the theoretical predictions, reduced variance leads to increased fitness, while increased variance leads to reduced fitness (Fig. 4a). These fitness changes are driven by the variance in the mother-daughter transitions (asymmetry), and less so by variance in growth when not dividing (Gr) or the variance in division probabilities (Div). We come to this conclusion as, when dropping growth variance and division variance, the population growth effect does not substantially change (compare Fig. 4a to Fig. 4b). Similarly, dropping the covariance term from the mother-daughter division asymmetry does not alter the effect of noise on population growth (compare Fig. 4b and 4c). We then ask whether mother daughter asymmetry variance in size or TF signal has the larger influence on population growth. The largest contribution of this altered variance effect comes from TF variance (compare Fig. 4d& Fig. 4e), where altering size variance show little effect on population growth. We caution that this last interpretation needs to be taken with consideration as variance is altered by multiplying with the same absolute value, but due to differences in the units (recall TF uses arbitrary units), the level of manipulation is not directly comparable. The stark differences in effect size suggest that variance in mother-daughter TF asymmetry is what influences population growth the strongest and where selection is expected to act most heavily on. Mother-mother division variance and mother-mother growth variance and covariance do not play an important role as does the covariance in mother-daughter transitions (Fig. S1). Note, all these latter (missing) variance effects likely do not cause a change in asymmetry, which strengthens our conclusion that mother-daughter TF asymmetry seems to be most sensitive and under strongest selection. Despite this expectation on increased selective strength acting against variance in TF division asymmetry, the observed variance in TF division asymmetry does not seem to be lower than that of growth asymmetry (Fig. 1c).

**Fig. 4:**
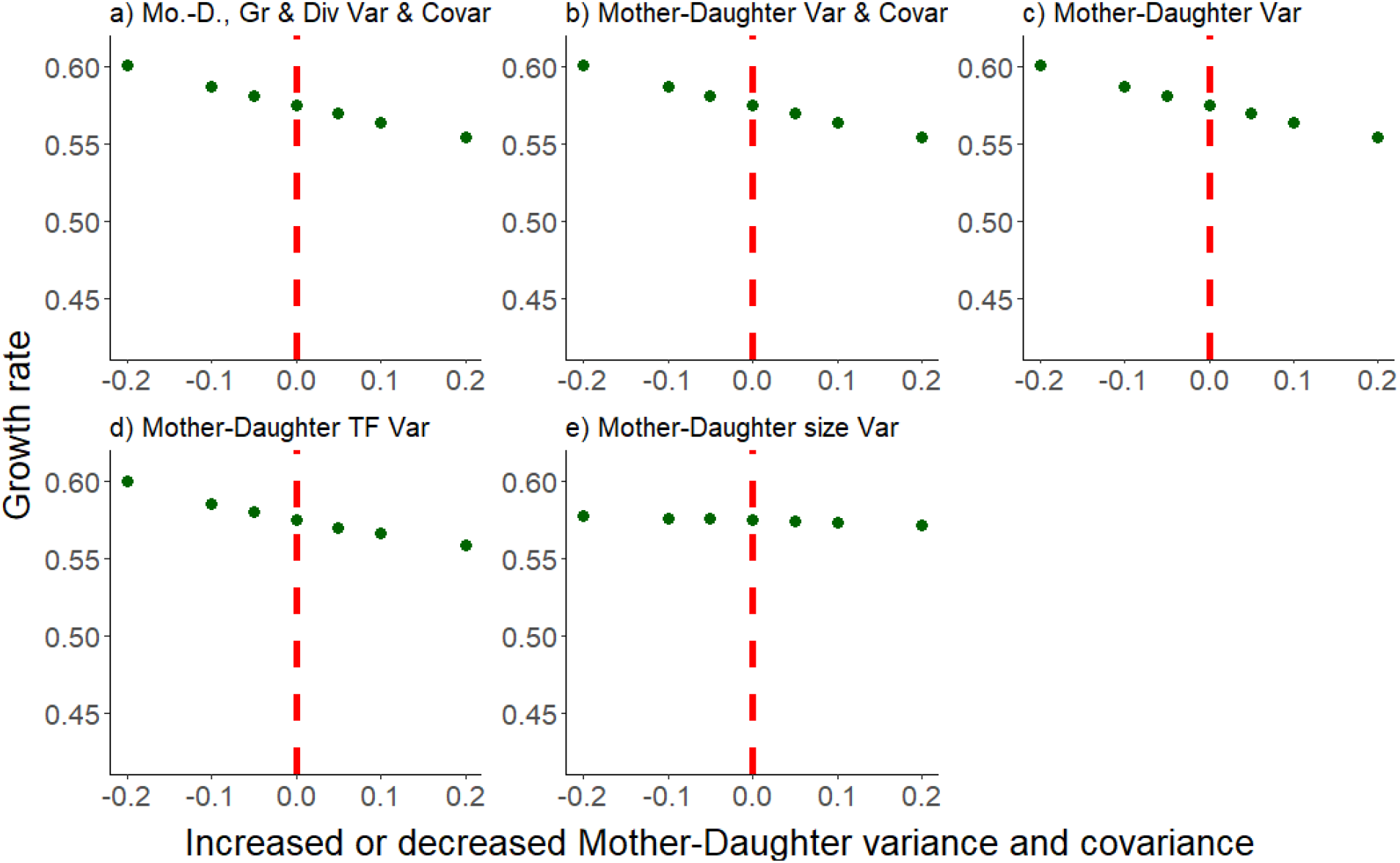
Effect on population growth of increased or reduced variance and covariance (noise) in mother-daughter transitions (Fertility matrix: *C*_1_(*z*^′^*l*^′^, *z, l*)), mother size transitions when not dividing (growth, Gr), and mother size transitions when dividing (Div). a) combined effect of increased or decreased noise of all three noise terms. b) effect of only the mother-daughter variance and covariance noise. c) effect of only mother-daughter variance (no covariance manipulated). d) effect of only mother-daughter TF variance (no covariance manipulated). e) effect of only mother-daughter size variance (no covariance manipulated). The red line represents the observed variance and covariance. Additional composition of division and growth variance and covariance contributions, as well as Mother-daughter covariance contributions, are shown in Fig. S1. These variables shown in Fig. S1 contribute only marginally to fitness if their noise levels are altered.

## Discussion

We ask two main questions: whether there is a division asymmetry that has likely evolved to an optimum, and how sensitive deviations from this optimum are, i.e. what type of cell intrinsic noise and variance in division asymmetry influences population growth the most. On the first question, we illustrate that the observed bacterial cell division asymmetry has likely evolved to optimize fitness. We derive this conclusion from the finding that reducing or increasing asymmetry leads to lowered population growth. Our study also illustrates a deeply integrated division process, as investigating only one of the two variables would have revealed different asymmetry optima compared to the one we find for integrating both variables (Fig. 3). The deeper integration might arise from underlying trade-offs, as it is well known that the transcription factor expression of *rpoS* negatively correlates with cell growth (5, 16). On the second question, we show that variance in mother-daughter TF asymmetry is most influential on population growth and hence should most heavily be selected against (14, 15), even though we do not see much reduced variance in asymmetry in that measure compared to other measures (Fig. 1c). These other variables such as variability in size or growth are of less importance for fitness, even though mean growth is closely related to fitness, but not to division asymmetry. Our approach of combining size and TF asymmetries in an advanced model that scales single-cell characteristics to population level dynamics allowed to reveal the deeper integration and decomposition of within and between mother and daughter cell stochasticity for fitness.

Previous models of aging and division asymmetry suggested that symmetrically dividing cells could only prevent population aging by a fine balance between cell growth (biomass gain), the accumulation of damage, and the dilution of damage by division (9, 20, 21). In contrast, asymmetry provides simple means that not only balance accumulation of damage by growth and division and damage related mortality, but also a partial rejuvenation of cells by asymmetric partition of damage (3, 4, 9, 20, 22–24). Models suggested that when damage accumulates with age, fitness—population growth— is maximised by a moderate asymmetric division of damage, while too little and too much asymmetry is sub-optimal (9). Unfortunately, to date, we have no causal mechanism that quantifies damage and explains observed mortality and demographic fates of cells (25). Our transcription factor signal dynamics of *rpoS*, does not serve as a damage marker, and the relationship between *rpoS* signal asymmetry and growth might be more complex than simple damage accumulation (5, 6). Our findings still align with asymmetric damage distribution as TF asymmetries show larger impact on fitness than size asymmetries. This in itself is interesting, as *rpoS* and growth are negatively correlated (5, 16). Only recently have systematic size differences at division in mother-daughter cell pairs been shown in *E. coli* bacteria (6, 8, 26), while growth difference between mother and daughter cells have long been established (1, 4, 25, 27, 28).

Asymmetries as source for increased fitness have also been investigated in the context of non-linearity of exponential cell growth and variance in division times (29). Similar findings revealed positive effects on population growth of variance in growth rates, size at division, and size at birth (30). These findings contrast with the general understanding that variance in fitness components should be selected against, where variance in vital rates reduce growth rates (18, 31–33). Our findings (Fig. 4) support that latter understanding when investigating the effect of increased noise (variance and covariance) in growth and TF signal. For most variance and covariances, we find small or negligible negative effects on fitness, as largely predicted (18, 31–33). Size and growth should be closely related to fitness, but we show that only variance in TF asymmetry negatively influenced fitness, which suggests that TF asymmetry variance might be under stronger selection than stochasticity in size and growth. This could be understood in light of our TF (*rpoS*) regulating about 10% of all *E. coli* genes (34). Size asymmetry might show multiple fitness optima (Fig. 3e), the reason for such different optima in size asymmetry are not easily understood. They might be driven by limits of the parameter space explored and that trade-offs between growth and *rpoS* expression cannot easily be swapped to a positive correlation, that reversed asymmetries would do. More detailed explorations would be required to understand the potential for various local optima and complex fitness landscapes. Further theoretical and empirical explorations could investigate error structures in more detail. Our model concerns gaussian distributed variance, but non-gaussian, skewed variance distributions, might lead to a stronger impact of altered noise, as effects such as Jensen’s inequality should gain relevance for fitness. Theoretically, such expectations have been formulated depending on autocorrelation in growth and perturbations (18, 35). Our expanded IPM model might provide one mode for such explorations of skewed distribution of noise.

Understanding optimal asymmetry and the selective forces acting on it, helps to explain basic biological processes such as aging in bacteria, and more general cellular aging that underlies all aging processes (25), it also enlarges understanding how heterogeneity among isoclonal individuals is generated, to the level that it might be the foundation of division of labour and a crucial condition for the evolution of multicellularity (36). The evolution of non-genetic non-environmental heterogeneity is yet surprisingly little understood, though there are some indications that such heterogeneity is not strongly selected against (37). On the one hand, our analyses supports that there is not much selection against noise, as increased noise does not influence fitness strongly. On the other hand, division asymmetry seems to be selected for, which leads to heterogeneity among individuals, that can be beneficial under varying environments as highlighted by bet-hedging arguments. Asymmetry can be similarly beneficial, for instance segregating (misfolded) protein aggregates asymmetrically—that traditionally have been associated with toxicity and accumulating damage—can increase survival under stressful conditions (38) and might help to buffer effects of antibiotic exposure (39, 40). Protein aggregates are also found to increase the fraction of antibiotic persister cells among antibiotic susceptible populations and a link to aging and persistence has been established (39–42). This non-genetic persistence is linked to the evolution of genetic antibiotic resistance, causing the failure of disease treatments (43–45). Such evolutionary properties of heterogeneity illustrate why we should shed more light on the processes generating heterogeneity, including the evolution and maintenance of asymmetry (37, 46). Exciting explorations lay ahead for such evolutionary understanding of asymmetry, stochasticity, and heterogeneity.

## Methods

### Data to parameterize IPM

To parameterize the model, we used single-cell data of *E. coli* strain MG1655 bacteria (5), collected using a mother machine microfluidic device (4, 28). The strain expressed a fluorescent marker under the rpoS promoter, which we label as transcription factor signal (TF) (Proenca *et al*. 2024). Cell length (growth), and TF signal were recorded at 4 minute intervals. Single-cell data on cell length was used to determine division events and division rates. Cells were grown in M9 media enriched with Casamino acids and glucose in the microfluidic device at 37°C. Each cell and its dividing offspring cells was tracked until it died, or was right-censored if the cell was washed out of the growth wells or was still alive at the end of the 36h experiment. We used a deep neural network based image analysis tool, DelTA (47) for which algorithms were trained on our data and in addition error detection and correction were made via R scripts, for details please see (5).

### Estimating Kernel parameters

To estimate parameters in a classical one-variable IPM, survival, *s*(*z*), is normally described as a binary logistic regression, as is the probability to reproduce, *p*_*b*_*b*(*z*). As mentioned, in binary fissioning bacteria, the number of offspring produced (*p*_*r*_ *b*(*z*)) can be set to one. The transition (growth) is usually computed as a linear regression with a variance distribution around. This variance considers that individuals with trait *z* at time *t* can transition to trait *z’* at time *t+1* given that they survive the interval (Ellner *et al*. 2016).

To estimate survival probabilities, *s*(*z, l*), for our two-variable IPM, we fitted a General Additive Model (GAM) with a binomial (logit link) error structure, with survival as response variable and the two variables *z* and *l* (i.e. cell length and TF signal intensity) as explanatory variables with an interactive smoothing term (Fig. 2a) (48). GAMs are penalized generalized linear models with local regression splines being used for the smoothing term. They are well suited for non-linear patterns. For multi-dimensional smooths, that we use for the two continuous variables, thin plate regression splines are applied (49).

To estimate reproduction (division probability) probabilities *p*_*b*_(*z, l*) we fitted a GAM with a binomial (logit link) error structure, for reproduction (division yes/no) as response variable, and the two variables *z* and *l* (i.e. cell length and TF signal intensity) as explanatory variables with an interactive smoothing term (see also Fig. 2b).

For the fertility matrix, **C**_1_(*z*^′^*l*^′^, *z, l*), we fitted a multivariate GAM where the size, *z*, and TF, *l*, of the mother cell before its division predicts the size, *z’*, and TF, *l’*, of the daughter cell after the cell division [R notation: gam(lDaughter ∼ s(lMother, zMother), zDaughter ∼ s(lMother, zMother)), family = mvn(d = 2))] (Fig. S2). These multivariate GAMs estimate the means, variance and covariance for both variables (size and *TF*) [R notation covariance matrix: solve(crossprod(multivariateGam))]. We assume normally distributed errors around the variance and covariance estimates, which allows estimating multinomial distribution for length and TF that can be used to parameterize the fertility matrix. Compared to previous IPMs with two variables where one variable is rather considered to be a covariate (13), we discretise the traits, and estimated for each *z, l* combination of a mother cell the probability distributions in producing a daughter cell with *z*^′^, *l*^′^ trait characteristics. Therefore, the matrix is inflated as for each *z, l* combination (Fig. S2). That means not only a single data point is estimated, but a matrix with the probabilities of producing a daughter cell with *z*^′^, *l*^′^ trait characteristics.

For time intervals, we assumed that a cell first needed to survive the interval and then evaluated whether it reproduced. Each of the *z, l* discrete value specific matrixes are then added and thereby collapsed to get again an original *C*_1_(*z*^′^, *l*^′^, *z, l*) matrix. To reduce matrix dimensions and simplify estimation we categorize the continuous variable *z, l* and *z*^′^, *l*^′^ combinations by defining ranges of *z* and *l*, based on 10 mesh points. We apply the midpoint rule to discretise cell size, *z*, and TF signal, *l*, with lower and upper limits being expanded to reduce edge effect biases (Ellner *et al*. 2016). This categorization is applied to all matrixes and regressions of the models.

For the mother cell growth-TF function, **G**(*z*^′^, *l*^′^, *z, l*), we estimated two separate multivariate GAMs, one for the event that a mother cell does not divide in the next time interval, and one for the event that a mother cell divides, which means that the mother cell at the next time interval is much reduced in size as a substantial fraction is partitioned to its daughter cell (Fig. S2). Similar to the mother-daughter transition **C**_1_(*z*^′^, *l*^′^, *z, l*), also for the growth-TF function, **G**(*z*^′^, *l*^′^, *z, l*), we estimate for each *z, l* combination the transition probability distribution to a *z*^′^, *l*^′^ mother cell, which results in for each *z, l* combination one matrix (in total 10×10 matrixes as we have 10*10 *z, l* categories). The matrix for each *z, l* combination, given that a mother cell does not divide, is then added to the matrix for each *z, l* combination, given that the mother cell divided. When adding *z, l* combination matrixes, we weighted each of these two matrixes based on the division probability *p*_*b*_(*z, l*) for such a *z, l* mother. We then multiplied the weighted and added *z, l* matrix that predicts the *z*^′^*l*^′^ cell in the next interval by the survival probability of *s*(*z, l*). These probability matrixes for each *z, l* combination predicting a *z*^′^, *l*^′^ cell in the next interval, were then summed to obtain one **G**(*z*^′^, *l*^′^, *z, l*) matrix that is of the initial size (10*10 mesh points).

### Perturbing mother-daughter asymmetries

We simulated altered asymmetries by taking the IPM parameterized with the observed data and perturbed the observed cell length of each mother cell after division by up to +/- 20%. At the same time, we perturbed observed cell length of the newly generated daughter cell by -/+ 20%. We did this in a reciprocal way so as to increase or decreased asymmetry; that is, if mother cells were enlarged, daughter cell’s size was reduced, or vice versa. We applied the same principle for TF signal, i.e. increasing and reducing mother-daughter cell asymmetries in TF signal after division. Perturbed asymmetries were explored for size and TF asymmetries independently (Fig. 3d & e), or jointly (Fig. 3a-c). Note, for these perturbations the original empirically recorded size and TF data was altered before it was log transformed, recall that the log transformation was primarily done for visualization and normalizing residual error distributions.

### Perturbing variances

In addition to exploring perturbations in mean mother-daughter asymmetry we also explore increased or decreased variance and covariance between size and TF within mother cells, and in mother-daughter cell asymmetries. That is, we increased and decreased noise, stochasticity, or uncertainty between i) mother-daughter cells (MD), ii) dividing mothers before and after division (MDiv), and iii) normal growing mothers that are not dividing in the next time step (MGr). To achieve this, we perturbed the variance-covariance matrix values that were estimated based on the model parameterized by the observed data by +/- 20%. To decompose contributions, we perturbed with three different variance-covariance matrices: the Mother-daughter variance-covariance matrix (MD), the mother-mother variance-covariance matrix when the mother divided (MDiv), or the mother-mother variance-covariance matrix when the mother did not divide (MGr). We further decomposed contribution of variance or covariance only, by perturbing single elements of the variance-covariance matrix, or multiple parameters at the same time. In that way we decomposed the variance effects of size and TF in combination or each of the factors in isolation. It also allowed to isolate covariance only effects. As mentioned, we can do that for each of the variance-covariance matrix (MD, MDiv, MGr). The simulations based on these perturbed matrixes provided population growth rates and hence fitness effects of increasing or decreasing variance and covariance.

## Acknowledgement

We thank Stephen Ellner for discussions on multivariate integral projection models, Jens Rolff for comments on an earlier draft of this study. We thank the HPC Service of FUB-IT, Freie Universität Berlin, for computing time. This work was supported by the Heisenberg Programme of the German Research Foundation grant 430170797 (U.K.S.), a Humboldt Research Fellowship, Alexander von Humboldt Foundation (A.M.P.), German Research Foundation grant 430174701 (M.T.); and a Marie Skłodowska--Curie Actions’ European Postdoctoral Fellowship grant 101069035 (M.T.).

## Supplemental information

**Fig. S1:**
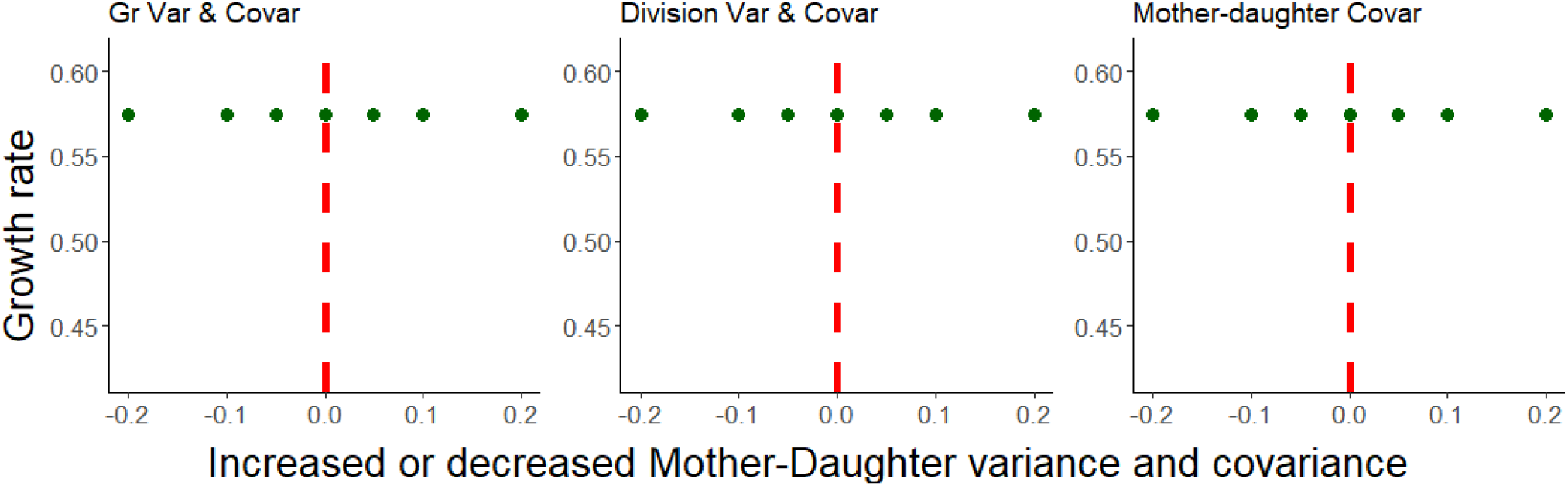
Effect on population growth of (a) increased variance and covariance (noise) in mother size (growth) transitions when not dividing, and (b) mother size transitions when dividing. (c) shows the covariance only effect of the mother-daughter transition. The red line marks the observed variance and covariance. Overall, altering variance and covariance in growth or division, or the covariance in division asymmetry among size and TF does not impact population growth, i.e. fitness.

**Fig. S2:**
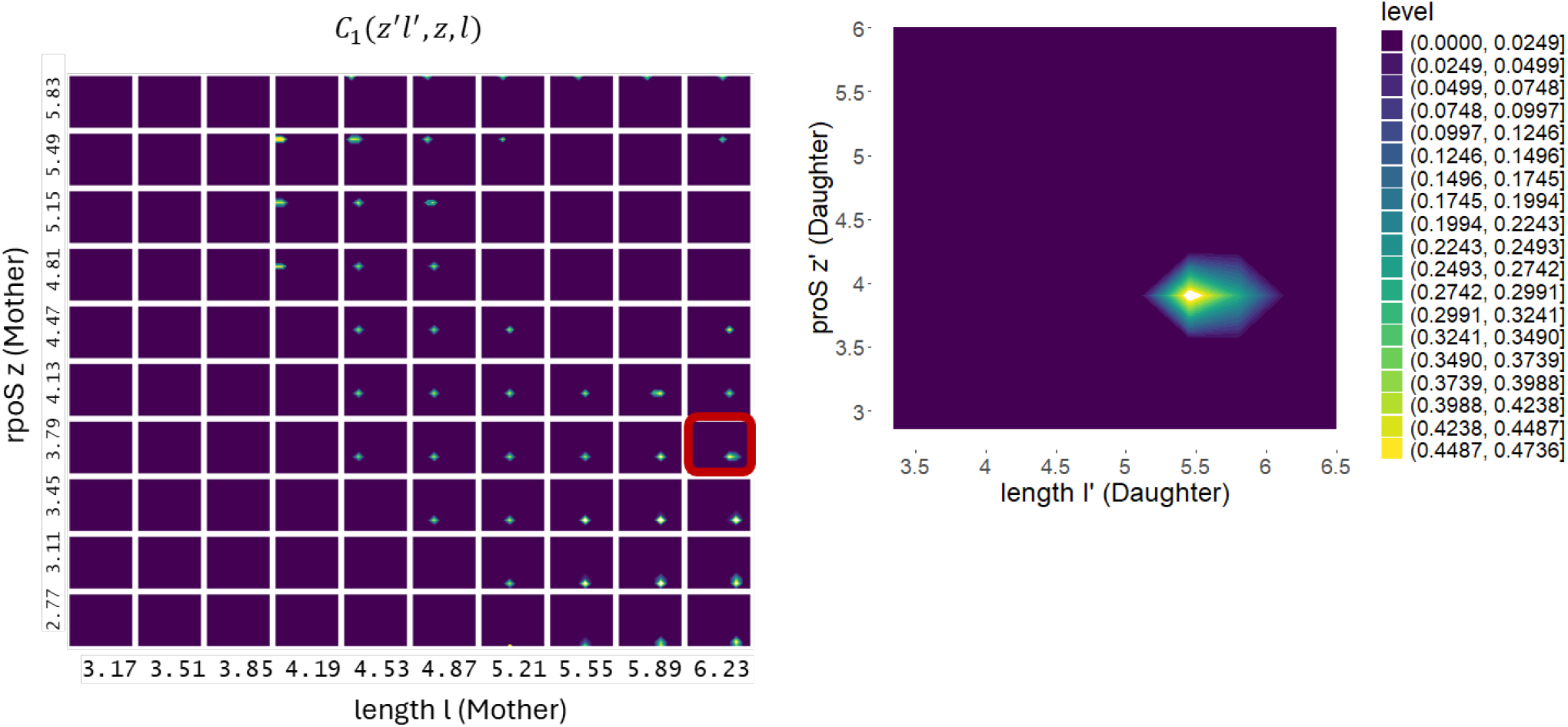
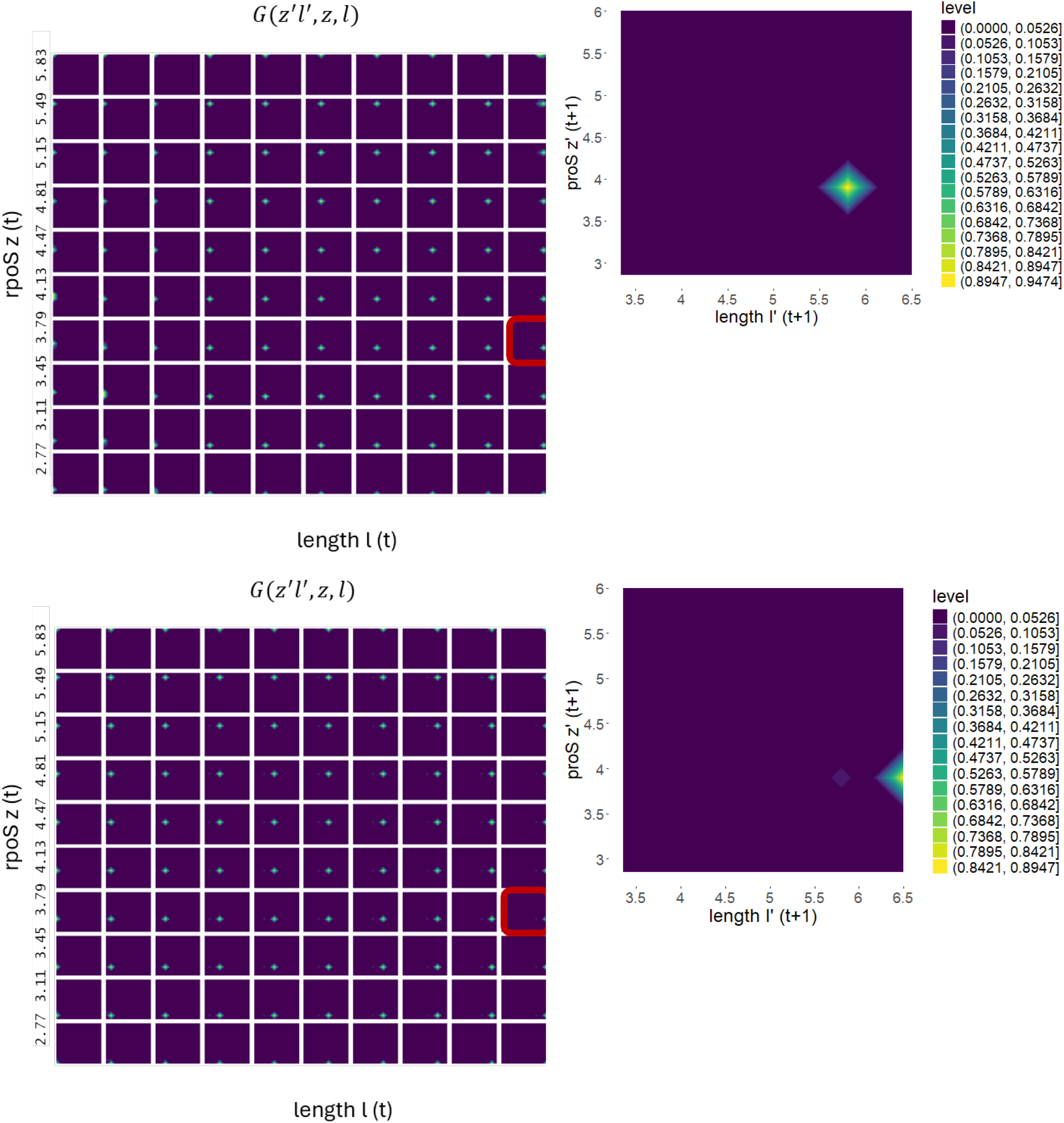
a) Mother-daughter transition *C*_1_(*z*^′^*l*^′^, *z, l*) distributions between time *t* and *t+1*. Mothers before dividing with length *l* and TF signal *z*, produce daughters of length *l’* and TF *z’* after division. b) Right panel example for a mother with 6.23 log length (midpoint) and 3.79 log TF (midpoint) produces daughters with length *l’* and TF *z’* with the shown probabilities. The example of b) is framed in red in a). Note, intensity of probability levels among each of the *z*,*l* combinations in a) are not equal. The difference in scaling among tiles of panel a) was chosen for better visibility. c) Mother-mother transition distributions *G*(*z*^′^*l*^′^, *z, l*) between time *t* and *t+1* when the mother is dividing c & d, or the mother is not dividing e & f. c) Mothers before dividing with length *l* and TF signal *z*, shrink in size (note the log scale) and are, after the division, of length *l’* and TF *z’* with the shown probabilities. d) Right panel example for a mother with 6.23 log length *l* (midpoint) and 3.79 log TF *z* (midpoint) is after the division of length *l’* and TF *z’* with the shown probabilities. The example of d) is framed in red in c). e) Mothers before dividing with length *l* and TF signal *z*, tend to grow in size (note the log scale) and are, after the division, of length *l’* and TF *z’* with the shown probabilities. f) Right panel example for a mother with 6.23 log length *l* (midpoint) and 3.79 log TF *z* (midpoint) is at the next time point of length *l’* and TF *z’* with the shown probabilities. The examples of d & f are framed in red in c) and e) respectively. Note, intensity of probability levels among each of the *z*,*l* combinations in c) are not equal. The difference in scaling among tiles of panel c) was chosen for better visibility. Note, intensity of probability levels among each of the *z*,*l* combinations within a and within e) are not equal. The difference in scaling among tiles of panel c & e) was chosen for better visibility.

**Fig. S3:**
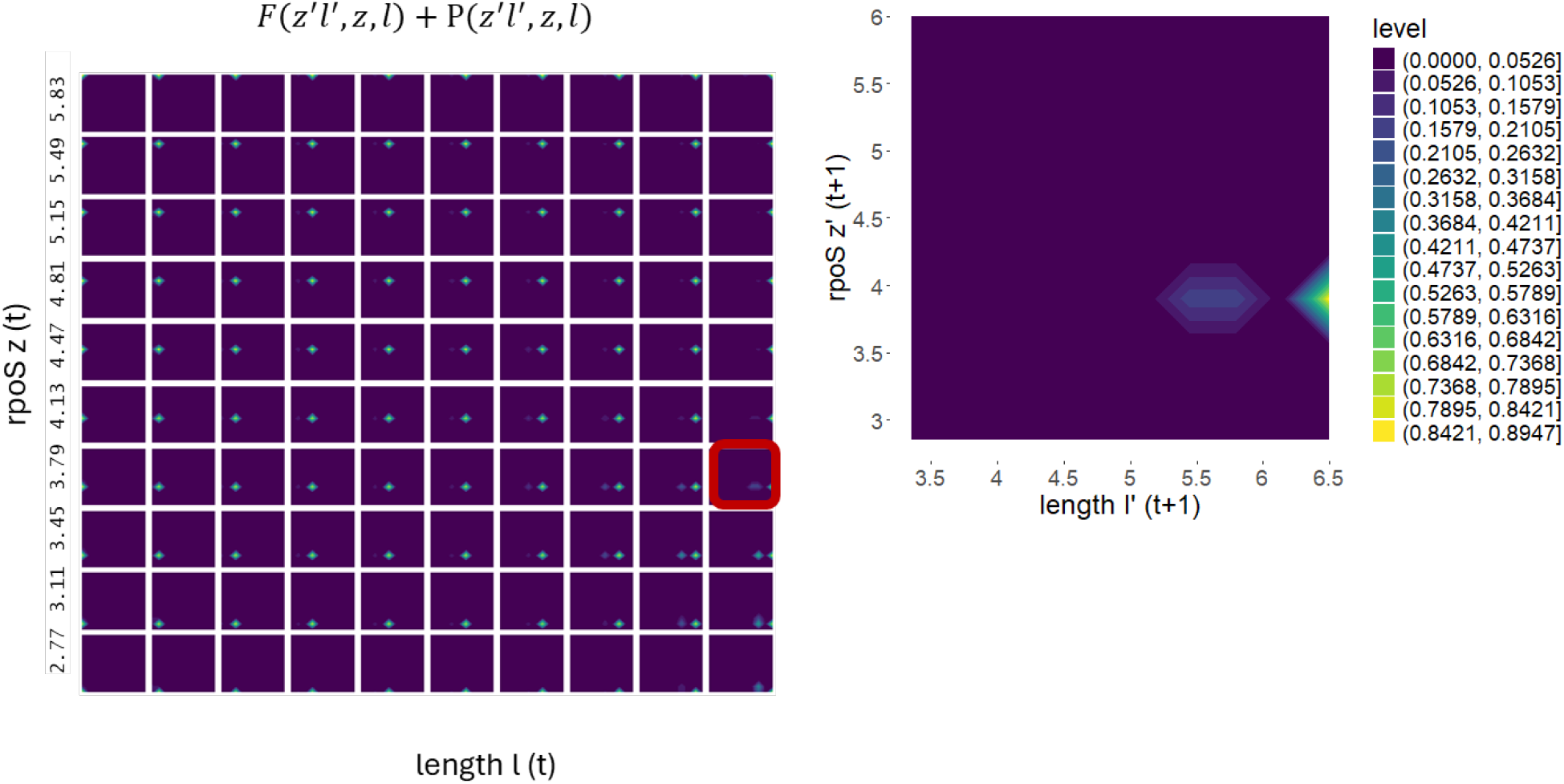
a) Combined kernel with fertility (mother-daughter) and mother-mother transition probability distributions **K**(*z*^′^, *l*^′^, *z, l*) = *p*(*z*^′^, l^′^, *z, l*) + *F*(*z*^′^, l^′^, *z, l*) that accounts for weighting of reproduction probabilities (see Fig. 2), growth transitions, and survival probabilities (Fig. S2) between time *t* and *t+1*. a) Mothers before dividing with length *l* and TF signal *z*, contribute to the next time (*t+1*) cell distribution of length *l’* and TF *z’* with the shown probabilities. b) Right panel example for a mother with 6.23 log length *l* (midpoint) and 3.79 log TF *z* (midpoint) and its contribution to the next time cells of length *l’* and TF *z’* with the shown probabilities. The example of b) is framed in red in a). In panel b) one can see the division probabilities (combined mothers when they divide and daughters, that are produced) as a lighter shading (smaller probabilities) at smaller sizes left of the growth of mother cells when not dividing.

